# Targeting DBS to the centrolateral thalamic nucleus improves movement in a lesion-based model of acquired cerebellar dystonia in mice

**DOI:** 10.1101/2024.05.21.595095

**Authors:** Megan X. Nguyen, Amanda M. Brown, Tao Lin, Roy V. Sillitoe, Jason S. Gill

**Affiliations:** Department of Pediatrics, Division of Neurology and Developmental Neuroscience, Baylor College of Medicine, Houston, TX, USA; Jan and Dan Duncan Neurological Research Institute at Texas Children’s Hospital, Houston, TX, USA; Department of Pathology & Immunology, Baylor College of Medicine, Houston, TX, USA; Department of Neuroscience, Baylor College of Medicine, Houston, TX, USA; Development, Disease Models & Therapeutics Graduate Program, Baylor College of Medicine, Houston, TX, USA

**Author notes:** **Co-corresponding authors:** Roy V. Sillitoe, Jason S. Gill.

**Keywords:** Dystonia, Cerebellum, Cerebellar Peduncle, Thalamus, Deep Brain Stimulation

## Abstract

Dystonia is the third most common movement disorder and an incapacitating co-morbidity in a variety of neurologic conditions. Dystonia can be caused by genetic, degenerative, idiopathic, and acquired etiologies, which are hypothesized to converge on a “dystonia network” consisting of the basal ganglia, thalamus, cerebellum, and cerebral cortex. In acquired dystonia, focal lesions to subcortical areas in the network – the basal ganglia, thalamus, and cerebellum – lead to a dystonia that can be difficult to manage with canonical treatments, including deep brain stimulation (DBS). While studies in animal models have begun to parse the contribution of individual nodes in the dystonia network, how acquired injury to the cerebellar outflow tracts instigates dystonia; and how network modulation interacts with symptom latency remain as unexplored questions. Here, we present an electrolytic lesioning paradigm that bilaterally targets the cerebellar outflow tracts. We found that lesioning these tracts, at the junction of the superior cerebellar peduncles and the medial and intermediate cerebellar nuclei, resulted in acute, severe dystonia. We observed that dystonia is reduced with one hour of DBS of the centrolateral thalamic nucleus, a first order node in the network downstream of the cerebellar nuclei. In contrast, one hour of stimulation at a second order node in the short latency, disynaptic projection from the cerebellar nuclei, the striatum, did not modulate the dystonia in the short-term. Our study introduces a robust paradigm for inducing acute, severe dystonia, and demonstrates that targeted modulation based on network principles powerfully rescues motor behavior. These data inspire the identification of therapeutic targets for difficult to manage acquired dystonia.

## Introduction

Dystonia is currently ranked as the third most common movement disorder[7, 23]. This high prevalence is in part due to the broad origins of the disease; dystonia may have underlying genetic, idiopathic, functional, neurodegenerative, and/or acquired causes[2]. However, regardless of the cause, dystonia describes the clinical phenomenon of abnormal extended contractions or co-contractions of muscle groups[2]. The underlying pathophysiology of different etiologies of dystonia varies widely, despite the convergence of the main motor phenotype. Importantly, the underlying etiology of dystonia has major implications for the success of response to treatment. Most amenable to treatment are the genetic and idiopathic dystonias, while acquired dystonia, which results from brain injury, is often more difficult to treat with existing approaches[11, 57, 85]. Of the dystonias that result from brain injury (including ischemic stroke, brain hemorrhage, and perinatal/*in utero* insults), cerebellar injury, specifically, has emerged as an important and powerful site of pathogenesis in both focal dystonia and more generalized dystonia[18, 43].

In clinical practice, focal lesions of the cerebellum, especially in adulthood, can lead to focal dystonia[19, 46]. Furthermore, in cerebral palsy (CP), the most common movement disorder in pediatric populations, developmental insults to the cerebellum may contribute to dystonia[66, 70, 78], which has been increasingly recognized as a key manifestation of the syndrome[24, 28]. The contribution of cerebellar injury to dystonia has been combined with neuroimaging investigations, model organism studies, and clinical therapeutics to delineate a “dystonia network” comprised of the basal ganglia, thalamus, sensorimotor cortex, and cerebellum[20, 40, 45]. Thus, understanding how acquired injury to cerebellar structures produces a dystonia with high morbidity that is difficult to treat has emerged as an important outstanding question.

The putative role of the cerebellum in dystonia, which is increasingly appreciated, reveals itself through the intimate positioning of the cerebellum at the nexus of sensorimotor integration[73]. The cerebellar cortex receives pan cortical afferent projections via pontocerebellar mossy fibers and accommodates ascending sensory information through olivocerebellar climbing fibers. Integration of the received sensorimotor information is then performed, predominantly, by granule cells and Purkinje cells in the cerebellar cortical circuit. The integrated sensorimotor model produced by the Purkinje cells is communicated to the cerebellar nuclei and from there to the thalamus using a major output pathway called the superior cerebellar peduncle (SCP)[30] (Figure 1A). From a neural network perspective, the SCP – connecting two crucial sensorimotor brain regions, the cerebellum and the thalamus – thus emerges as a key focus for investigating the neural mechanisms in dystonia pathogenesis.

**Figure 1.**
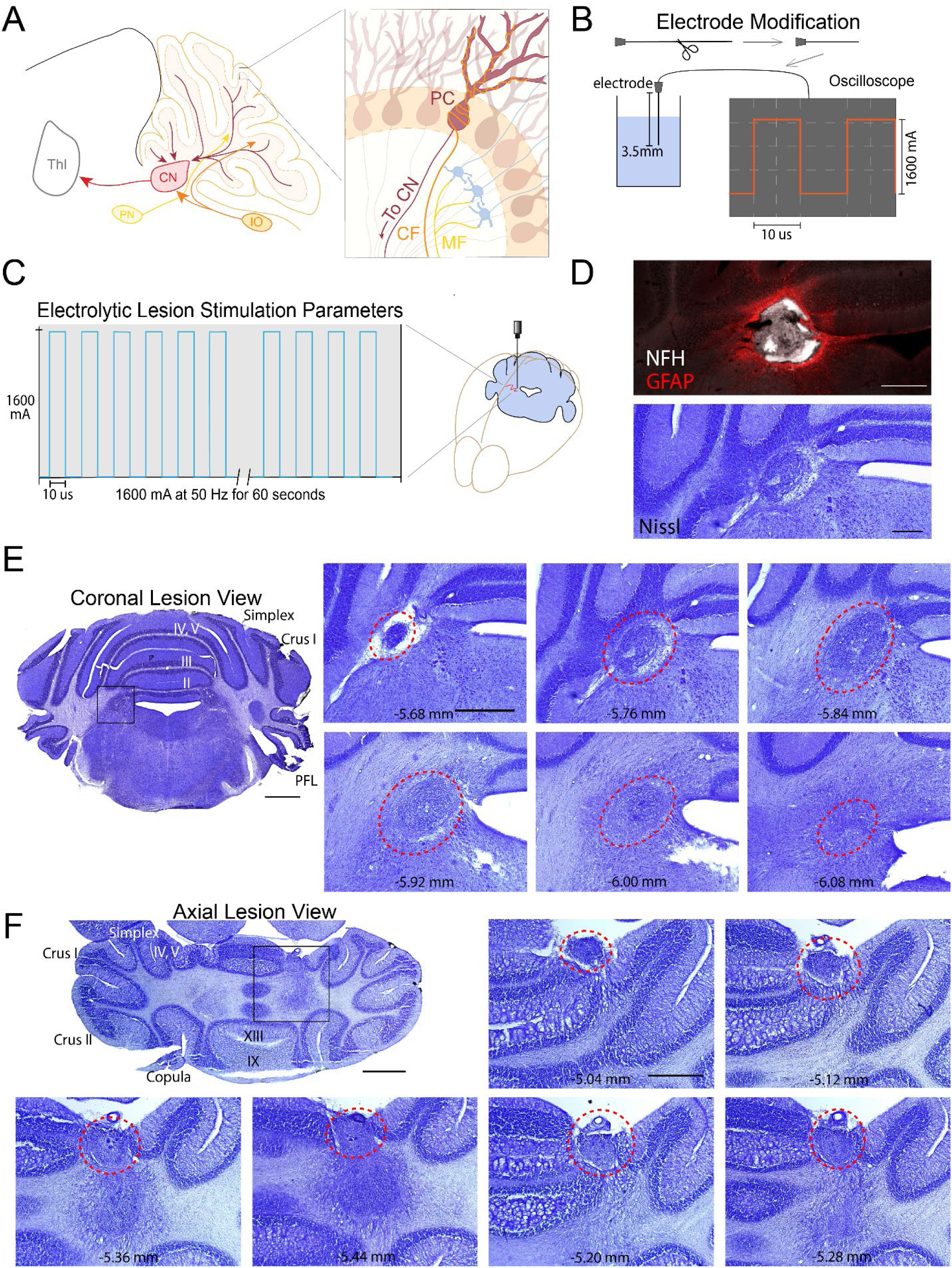
The cerebellar network and electrolytic lesion paradigm. A simplified diagram showing example primary afferent projections to the cerebellar cortex, such as the pontine nuclei (PN), which transmit descending cerebral cortical information to the cerebellar cortex via mossy fibers (MF) (A). Inferior olivary neurons (IO) project exclusively to the cerebellar cortex via climbing fibers (CF) (A). Purkinje cells (PC) then assimilate all cerebellar cortical computation and form the sole output neurotransmission of the cerebellar cortex, projecting mainly to the cerebellar nuclei (CN) (A). The CN project via the superior cerebellar peduncle (Red) to the thalamus (Thl) (A), among other regions. For electrolytic lesioning, tungsten electrodes are cut at 3.5mm, and then tested using an oscilloscope to verify capability to deliver a 50 Hz 1600 mA charge (B). The modified electrodes are then stereotactically inserted to a target location in the brain and the 1600 mA, 50 HZ charge is delivered for 60 seconds (C). Nissl stain at site of lesion reveals cystic changes at target site (D, lower panel). Immunohistochemistry with GFAP (red) and neurofilament heavy chain (NFH; white) reveal gliosis and neuron degeneration at the lesion site, respectively (D, upper panel; scale bar = 250 um). Lesion volume in the coronal (E) and axial (F) plane are shown (lobules are as noted, PFL=paraflocculus). Serial sections are shown through the lesion with distance from bregma indicated in the inset. Red circles represent the same lesion across sections. Lesion volume is roughly a sphere with diameter of 0.4 mm. All sections used in the imaging were obtained from animals 2 weeks after lesioning (scale: gross image = 1 mm; insets = 500 um).

The contribution of afferent cerebellar projections and cerebellar cortical manipulations to dystonia has been supported by a variety of genetic, structural, and functional manipulations performed in rodent models, which are uniquely suited for dystonia investigations given the evolutionary conservation of the cerebellar circuit across mammalian species[9, 30]. Mouse models of the monogenic dystonia, Dyt1, have revealed dysfunction and degeneration of Purkinje cells and cerebellar nuclei neurons[26, 49, 87]. Building upon these findings, the monogenic rat dystonia model, *dt*, implicated olivocerebellar dysfunction in dystonia through electrophysiological analysis of Purkinje cell and cerebellar nuclei neuron firing[47]. Furthermore, in Dyt12, a rapid onset dystonia parkinsonism syndrome, pharmacological manipulation of Purkinje cells has been shown to be sufficient to induce dystonia[14, 25]. Moving on from Mendelian models of dystonia, pharmacological neuromodulation/lesioning of the cerebellar vermis through kainic acid-induced glutamatergic excitotoxicity has also been shown to be sufficient to produce dystonia[63]. The fine scale manipulation of cerebellar circuitry through conditional genetic approaches has compellingly revealed dystonia arising from aberrant function of the cerebellar circuit [9, 31, 48, 83]. This series of papers revealed that silencing of afferent inferior olivary projections produces dystonia early in development and persisting into adulthood[9, 31, 48, 83]. In this model, loss of input from the olivary projections leads to altered Purkinje cell firing and subsequent disruption of normal cerebellar nuclear firing patterns[83]. Together, these genetic, structural, and functional manipulations have shown that cerebellar circuit dysfunction – dysfunction of afferent signaling at the input stage of the circuit and pre cerebellar nuclear processing at the output – is sufficient to produce dystonia.

Canonically, dystonia that is refractory to pharmacological therapy is treated with deep brain stimulation (DBS)[44, 77]. In genetic and idiopathic dystonia, DBS that is targeted to the globus pallidus interna (GPi) or the subthalamic nucleus (STN) is so efficacious that it can be considered a first line therapeutic option[36, 39]. In acquired dystonia, on the other hand, DBS targeting of these structures shows reduced efficacy in treating dystonia, and those that do respond have more persistent disability than their counterparts with genetic dystonia[52], pointing to differences in underlying network dysfunction[42]. This relatively poor response to GPi and STN targeting in acquired dystonia has led to the need for non-canonical targeting of other subcortical structures in the dystonia network – including the thalamus, SCP, cerebellar nuclei, and the cerebellar cortex – with varying results[38, 50, 51, 56, 60, 76, 86]. A focused interrogation of how deleterious (lesions) and therapeutic (DBS) perturbations in the dystonia network interact with dystonic phenotypes may help to parse these disparate observations.

Integrity of the SCP and cerebellothalamocortical connectivity may reflect the penetrance of dystonic phenotypes in patients with genetic dystonia[5]. We previously reported reduction of dystonic phenotypes through neuromodulation of the cerebellar nuclei, the final output cells of the cerebellum, in the olivocerebellar model of cerebellar cortical dystonia[83]. Subsequently, both pharmacologic silencing and DBS of the cerebellar nuclei were performed and found to reduce the dystonic phenotype in these mice[83]. In humans, iatrogenic lesioning of efferent cerebellar nuclear projections emanating from the cerebellar nuclei results in motor and cognitive-affective deficits in proportion to the extent of efferent tract damage[3]. The SCP itself is a key tract involved in motor adaptation[6]. Previously, studies involving lesions of the SCP or cerebellar nuclei (fastigial) have been performed, reporting degenerative changes in connected structures[15], and abnormalities in motor, cognitive, and social behavior[1, 33, 34]. Given the crucial position of the SCP in connecting two key loci required for sensorimotor integration, the cerebellum and thalamus, in the dystonia network, we posit that structural manipulation of this link may yield important insights into dystonia pathogenesis and treatment[56, 64, 73].

To investigate how disruption of cerebellar connectivity directly impacts motor behavior in mice, we developed a novel electrolytic lesion paradigm using modified metal electrodes. We targeted the SCPs bilaterally as they emerge from the medial and intermediate cerebellar nuclei, a target derived from clinical studies implicating damage at this site as a crucial determinant of broad cerebellar mediated behavioral dysfunction[3](Figure 1). We directed DBS to key nodes in the dystonia network to better understand how neuromodulation affects dystonia resulting from cerebellar outflow lesions. We find that lesions to the cerebellar outflow tracts are sufficient to produce a dystonia and that thalamic DBS reduces the severity of the resulting dystonia.

## Methods

### Animals

All mice used in the described experiments were housed in a Level 3, AALAS-certified facility. The Baylor College of Medicine (BCM) Institutional Animal Care and Use Committee (IACUC) reviewed and approved all studies performed using mice. All animals used for the included experiments were C57 Black 6J mice (Jax: 000664). Mice were purchased from JAX and then bred and maintained at BCM. All experiments included mice of both sexes. The day of birth was considered postnatal day 0 (P0). All mice tested were between P28 and P42 (4-6 weeks of age). The mice were housed under a 14 hrs/10 hrs light/dark cycle, with daily temperatures held between 68 and 72 °F, and humidity kept between 30 and 70%.

### Tissue processing

We collected brain tissue for analyses as previously described[80]. Mice were anesthetized with 2, 2, 2-tribromoethanol (Avertin). Deep pressure toe-pinch was used to ensure adequate anesthesia. Mice were then perfused through the left ventricle of the heart with 0.1 M phosphate buffered saline (PBS, pH 7.4) until clear PBS flowed from a nick in the right atrium. This was followed by perfusion with 4% paraformaldehyde (PFA) diluted in PBS for fixation. The brains from the perfused mice were dissected and postfixed for 24–48 hrs in 4% PFA at 4 °C and then cryoprotected stepwise in PBS-buffered sucrose solutions (15 and 30% at 4 °C for 24-48 hrs each). Next, the tissue was frozen in an optimal cutting temperature (OCT) solution and stored at −80 °C. Serial 40-µm-thick tissue sections of the orientation indicated in the results description were obtained using a cryostat, and then collected and processed free-floating in PBS.

### Nissl staining

Tissue sections were mounted and dried overnight on glass slides. After drying, the mounted sections were immersed in 100% xylene twice for five minutes. They were then rehydrated through a series of four immersions in: 100% ethanol, 95% ethanol, 70% ethanol, and then water; each step for 2 min. Subsequently, the sections were immersed in cresyl violet solution until the appropriate staining was achieved, which was about two minutes. The sections were then dehydrated in 70%, 95%, 100% ethanol followed by immersion in 100% xylene, for one minute each. The slide processing was then completed immediately by mounting with coverslips using Cytoseal XYL mounting media (Thermo Scientific, Waltham, MA, USA, #22-050-262). The cover slipped slides were left to dry overnight in a fume hood prior to imaging each tissue section.

### Immunostaining

Perfusion and tissue fixation were performed as previously described [74]. Mice were anesthetized and fixative was provided as described above in “tissue processing”. Samples were cut on a cryostat with a thickness of 40 µm and then processed as free-floating frozen tissue sections as described previously[75, 82]. After staining, the tissue sections were placed on electrostatically coated glass slides and allowed to dry. For detect glial infiltration at the lesion site, anti-rabbit GFAP (DAKO Catalog #Z0334) was used at a 1:500 dilution. For neurons, mouse neurofilament heavy chain (NFH; Covance Catalog #PCK-592P) was used at a 1:500 dilution. Brain tissue sections were incubated overnight for ∼16 hrs. Tissue sections were washed 3–4 times with 0.1 M PBS for 5 min. The tissue sections were then incubated for 2 hrs with immunoglobins conjugated to Alexa 488 or 555 fluorophores (Invitrogen Molecular Probes Inc., Eugene, OR, USA #A-21202, #A-31572) diluted 1:1500 in 10% NGS blocking solution. Tissues were then rinsed an additional 3–4 times with 0.1 M PBS. The fluorescently stained sections were mounted onto glass slides and then cover slipped with a DAPI-based mounting medium (Vectashield Anti-Fade Mounting Medium with DAPI #H-1200).

### Microscopy and image processing

Photomicrographs of sections were acquired with a Leica DM4000 B LED microscope and DPX365FX camera or using a Zeiss Axio Imager. M2 microscope with an AxioCam MRc5 camera. Whole section images were digitally stitched using the Adobe Photoshop Photomerge function. Color, brightness, and balance were adjusted using ImageJ software, and image processing for figures was completed using Adobe Illustrator.

### Surgical procedures

The surgical approach used in the current experiments was derived from methodology previously described[55]. All surgical techniques used for the described studies began with the same induction protocol: buprenorphine (slow release, 1.0 mg/kg sub-cutaneous (SC)) and meloxicam (5.0 mg/kg SC) were given for pre-surgical analgesia with ongoing application provided for post-operative recovery. Induction of anesthesia was achieved with 3% isoflurane gas and verified using deep toe pressure. Anesthesia was maintained during surgery using 2% isoflurane gas administered through a mouthpiece on the surgical stage. Surgery was performed on a stereotaxic platform (David Kopf Instruments, Tujenga, CA, USA) using sterile technique from start to finish. For surgeries, an incision was made to expose the skull anterior to bregma and posterior to expose the insertion of the muscles on the occipital skull. Following surgery, the mice were placed in a pre-heated warming chamber for monitoring and recovery.

### Lesion paradigm

Electrodes were prepared for lesion induction as follows. Tungsten electrodes (Thomas Recording, Germany) were cut to 3.5 mm (Figure 1). To ensure that the electrode would deliver the desired charge, each electrode was attached to a Multichannel Systems pulse generator (AMPI, Jerusalem, Israel) programmed to deliver 1600 mA square pulses at 50 Hz. The end of each electrode was then placed in saline solution. Current amplitude, pulse width, and frequency was confirmed by an oscilloscope (Tektronix TBS 1062) using two leads spanning a 10 kΩ resistor. The resistor was placed with one end attached to a ground wire and the other submerged in the saline solution. Electrodes that delivered charge mirroring that programmed into the Multichannel Systems pulse generator as reflected by the oscilloscope readings were then selected for use in the lesion induction.

For the lesion and sham lesion surgeries, mice were anesthetized, and the skull was exposed as described above. A craniotomy site was prepared bilaterally at -5.7 mm A/P (from bregma) and +/-1.2 mm M/L to expose the brain parenchyma overlaying the SCP at it emergence anterior to the fastigial (medial) and interposed (middle) cerebellar nuclei based on Paxinos and Franklin [62]. A prepared electrode was then inserted into the craniotomy site and lowered -2.4 mm (0 mm was considered to be at the point the electrode contacted the exposed dura). For lesioning, square pulses of 1600 mA were delivered at 50 Hz for a duration of 60 s. For sham lesions, the electrode was inserted and left at the target depth for 60 s without delivery of the electrical charge. For either lesion or sham animals the same procedure was then repeated on the contralateral side of the brain.

### DBS

Twisted bipolar tungsten electrodes with a width of 0.127 mm and a length of 3.5 mm were purchased from PlasticsOne. Two electrodes were spaced 3.1 mm apart to bilaterally target the centrolateral thalamic nuclei or 3.5 mm apart to target the dorsal striatum. They were then fixed together using Bondic, a UV light-activated bonding agent (Bondic, Niagara Falls, NY, USA). As described above, the mice were then deeply anesthetized with isoflurane so that the DBS electrodes could be inserted into the target region based on stereotactic coordinates derived from Paxinos and Franklin (2001): for centrolateral thalamic nuclei -2.06 mm (anterior–posterior; bregma = 0 mm), +/-1.55 mm (medial–lateral), and -2.55 mm (dorsal–ventral); for dorsal striatum 0 mm (anterior–posterior; bregma = 0 mm), +/-1.75 mm (medial–lateral), and -2.0 mm (dorsal– ventral)[62]. Electrodes were secured to the head with C&B Metabond (Parkell, Inc., Edgewood, NY, USA, SKU: S380) and Teets ‘Cold Cure’ Dental Cement (A-M Systems, LLC, Carlsborg, WA, USA, Catalog #525000 and #526000).

### Electromyography (EMG)

To quantify *in vivo* muscle activity responses to DBS, EMG was used to monitor muscle contractions and the approach was adapted from previously described protocols[68]. EMG headmounts were constructed using silver wire electrodes (A-M Systems, Sequim, WA, USA; #785500) soldered to a connector for a detachable preamplifier (Pinnacle Technology, Inc, Lawrence, KS, USA; #8406). Subsequently, under deep anesthesia, the trapezius muscle was exposed bilaterally with preserved continuity from the craniotomy site created for lesion induction and DBS placement. Two silver wires (∼2.5mm apart) were inserted into each of the left and right trapezius muscles. Prior to insertion, the last millimeter of wire was folded back so that it would hook into the muscle, preventing dislodgment. A ground wire was then implanted into the subcutaneous tissue overlaying the muscle and the EMG head mount was secured to the head with C&B Metabond (Parkell, Inc., Edgewood, NY, USA, SKU: S380) and Dental Cement (A-M Systems, LLC, Carlsborg, WA, USA, Catalog #525000 and #526000).

### In vivo electrophysiology

Electrophysiological recordings of Purkinje cells were obtained from awake behaving mice according to previously described publications[10, 81]. Mice were surgically prepared for craniotomy as described above. For *in vivo* electrophysiology access, an ovoid craniotomy roughly 2 mm in diameter was created and centered roughly at -6.5 mm posterior to bregma and 1.3 mm lateral. For mice in which recordings were to be performed, it was ensured that the craniotomy encompassed both the coordinates required for the lesioning experiment and the recording site. The craniotomy was then protected using a custom-built 3D-printed chamber and filled with antibiotic ointment. To stabilize the mouse’s head during recordings a custom headplate was affixed over Bregma. The craniotomy chamber and headplate were anchored by insertion of a skull screw placed into an unused region of skull. The implanted items were secured using Metabond Adhesive Cement (Parkell, Edgewood, NY, USA) followed by dental cement (A-M Systems, LLC, Carlsborg, WA, USA, Catalog #525000 and #526000).

### Analysis of in vivo electrophysiology

Analysis was conducted as previously described[32]. Electrophysiological recordings were spike sorted in Spike2. Purkinje cell simple spikes and complex spikes were sorted out separately. Complex spikes were defined by their large action potential waveform that is typically followed by 3–5 smaller spikelets. The presence of complex spikes was used to define a recorded cell as a Purkinje cell for inclusion in our analysis. As a result, in the final analyses, Purkinje cells with clear complex spikes, high signal-to-noise ratio, and 60 s or longer duration of recording were included. We defined “firing rate” as the average number of spikes per second over the duration of a reliable recording. “CV” was defined as the standard deviation of the interspike intervals divided by the mean of the interspike intervals. For “CV2”, the following formula was used: 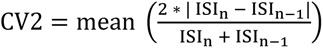, where ISI represents a single interspike interval in the spike train[37].

### Behavioral evaluation (mouse dystonic behavior)

The dystonia rating scale was adapted from methods described previously[63]. After completion of the surgery, mice were placed in a warmed chamber for monitoring and recovery. At 30, 60, and 120 mins after removal of the inhaled anesthesia, mice were examined and recorded for 60 s. Mice were scored based on the dystonia rating scale as described in Figure 3. A given mouse was scored for the *best* behavioral performance on the scale. For instance, if the mouse was in a fixed immobile posture for a significant duration of the study but was able to ambulate, although for only a short duration, the mouse would be given a score of 4.

### Statistical analysis

All statistical analyses and data presentation were prepared in GraphPad Prism (V10.1.0). As no differences based on sex were noted on initial analysis, mice from both sexes were combined across experiments and for statistical analyses (total mice n=27; female n=9, male n=18; Dystonia rating scale: lesion group: female=1, male=7:: sham group female=1, male=4; Deep brain stimulation: lesion with thalamic DBS: female=1, male=7:: sham with thalamic DBS female=1 male=4:: basal ganglia DBS: female=4, male=3; electrophysiology: lesion group: females=2, males=1:: sham group female=2, males=1). All *post hoc* pairwise comparisons were made with Bonferroni corrections. Paired two tailed t-tests were used for comparison of dystonia rating scale scores and EMG burst duration for thalamic DBS with sham surgery, thalamic DBS with lesion, and striatal DBS with lesion. One way ANOVA was used to analyze the difference between the percent change in burst duration between the three conditions (thalamic DBS with sham lesion; thalamic DBS with lesion; striatal DBS with lesion). As the striatal DBS animals did not show significant differences compared to the thalamic sham DBS group, striatal DBS on sham lesion animals was an omitted condition to minimize the number of animals undergoing the invasive surgical procedure.

## Results

### Devising an electrolytic lesion paradigm to induce focal lesions in the output pathways of the mouse cerebellar system

Previous mouse models using non-mendelian approaches to evaluate dystonia etiologies in mice have relied on conditional genetics or pharmacologic lesions[9, 14, 25, 31, 63, 83]. In patients, however, there are a variety of acquired brain insults that lead to focal lesions of the brain parenchyma, which occur in and around the cerebellar circuit that produce dystonia[18, 43]. Knowing that a variety of manipulations of afferent signaling to the cerebellar cortex, and disruption of cerebellar cortical function both lead to dystonia, we sought to devise a paradigm that would allow for the reliable induction of highly focal lesions to test whether disruption of cerebellar efferent signaling would also produce dystonia. Previous groups have used monopolar electrodes to induce lesions of the fastigial (medial) nucleus in rats[1, 33, 34]. Therefore, using tungsten electrodes, we sought to deliver a charge of 1600 mA at 50 Hz using a pulse generator attached to the electrode.

To verify sufficient charge delivery, we used an oscilloscope to measure the current produced by the electrodes. We modified the electrode by cutting it to 3.5 mm to achieve the desired depth and resistance (Figure 1B). Upon testing the modified electrode, the oscilloscope registered the expected charge delivery at the programmed voltage and frequency, with fidelity to the programmed waveform (Figure 1B). We then performed a surgery using the modified electrode, either by inserting the electrode and then removing it after 60 s (sham) or by delivering a charge of 1600 mA for 60 s at 50 Hz (Figure 1C). After surgery, we waited two weeks for lesion evolution prior to characterizing the impact of the resulting lesion. After taking the tissue from the animals, we used a Nissl stain to reveal the presence of hypercellular infiltrate surrounding a core consisting of cystic degenerative changes to the brain parenchyma (Figure 1D, see loss of neuronal morphology with NFH at lesion core). Immunostaining for GFAP confirmed that the hypercellularity consisted of glial cells, consistent with gliosis at the site of injury (Figure 1D).

Next, we used histologically stained serial tissue sections to examine the lesion size (Figure 1E, F). Using serial sections across two planes (coronal and axial), we found that along the anterior-posterior axis (using coronal sections), the lesion depth was approximately 0.4 mm (Figure 1E); this matched the lesion volume of approximately 0.4 mm along the dorsal-ventral axis (using axial sections) (Figure 1F). After optimizing the stereotactic targeting of the SCP, we noted that across animals with a successful and accurate lesion induction, the lesion characteristics at the level of tissue anatomy remained consistent with that shown in Figure 1D-F.

### Focal lesions in the cerebellar outflow tracts induce neuropathological features

The majority of efferent cerebellar throughput – ascending to the cerebral cortex and descending to the peripheral nervous system – is thought to pass through the SCP primarily from the cerebellar nuclei (Figure 1A). The efferent projections of the cerebellar nuclei through the SCP have been termed the cerebellar outflow tract[3]. In humans, acute lesioning of the cerebellar outflow tract leads to a constellation of symptoms termed the cerebellar cognitive affective syndrome, the severity of which occurs in direct proportion to the percent damage to the tract bilaterally[3, 35]. In rats, lesioning of the medial (fastigial) cerebellar nuclei has also been associated with a constellation of behavioral deficits[1, 33, 34]. In mice, the sequelae of lesioning of the cerebellar outflow tracts have not been examined. Taking advantage of the high degree of anatomical conservation of the cerebellar circuit between humans and mice, we derived the cerebellar outflow tract location from Albazron et al to target the SCP as it emerges from the medial and interposed cerebellar nuclei using stereotactic coordinates based on Paxinos and Franklin[3, 62]. Our stereotactic approach was able to reliably produce lesions with a core of cystic degeneration just anterior to the medial cerebellar nuclei using coordinates, from bregma, of -5.7mm posterior, +/-1.2 mm bilaterally, and at a depth of 2.4mm dorsal to the brain surface (Figure 2A, Figure 2C).

**Figure 2.**
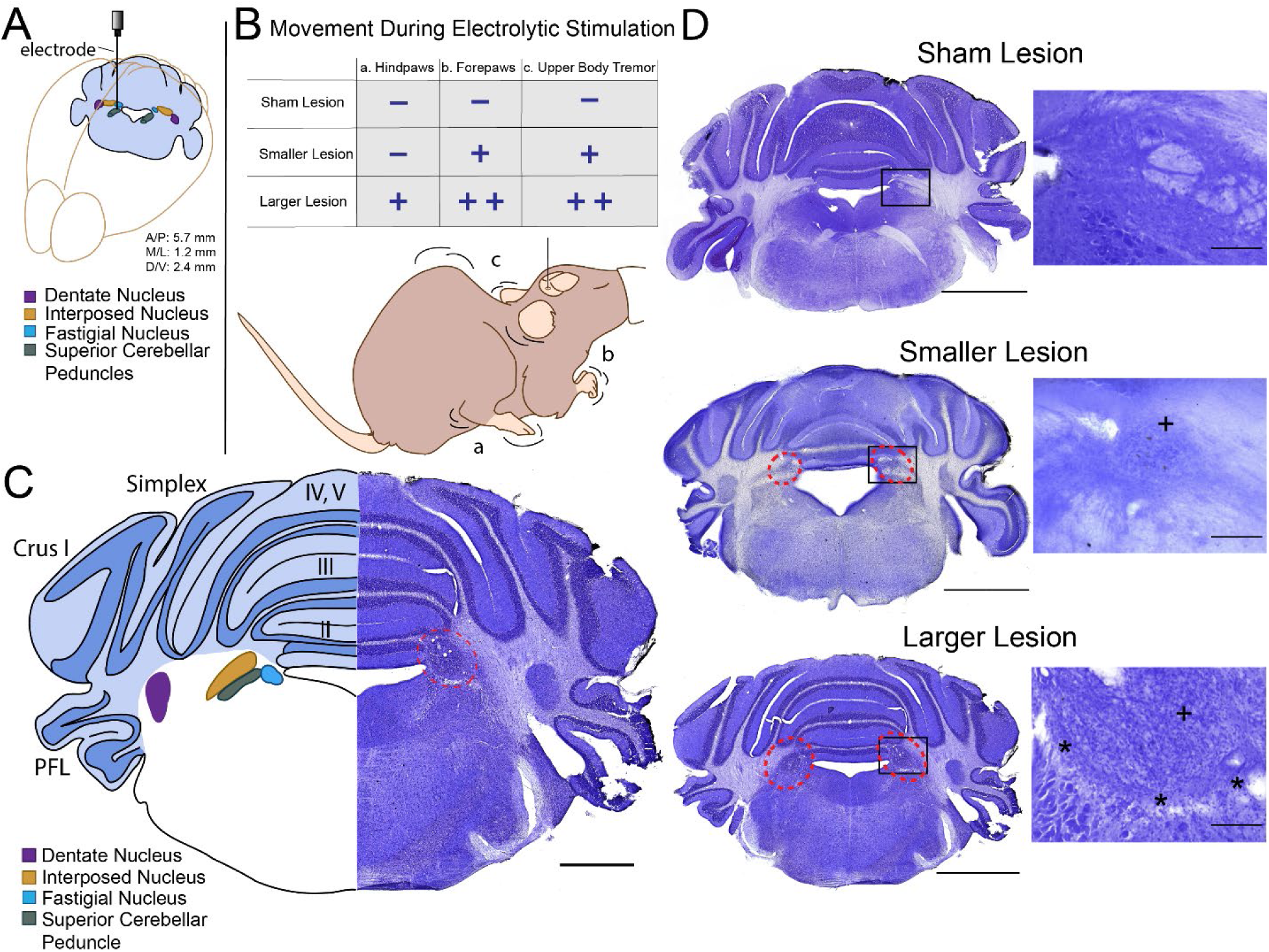
Electrolytic lesioning of the cerebellar outflow tract. Lesions were targeted to the superior cerebellar peduncle (SCP) as it emerges from fastigial (FN) and interposed (IN) nuclei (A, blue represents a section at the approximate coronal plane at which the lesion was made). This site was chosen based on the work of Albazron et al[3]. During lesioning, three classes of movements were noted (B). With sham stimulation, no movements were noted; with lesions that were smaller than expected there was mild forepaw movement and tremor of the upper body; with larger lesions hind paw movement was noted, more prominent movement of the forepaws, and tremor with contraction of trunk was seen (B). Lesion location (C, right) was centered on the site of convergence of the SCP, FN, and IN (C, left; lobules are as noted, PFL=paraflocculus; scale bar = 1mm). Sham lesions reveal no change to the brain parenchyma (D, upper); smaller lesions reveal gliosis (increased cell density indicated within red dotted lines on D middle, and + on inset image) without cystic changes (D, middle); larger lesions reveal more extensive gliosis (increased cell density indicated within red dotted lines on D bottom left, and + on inset image) with cystic changes (asterisks on inset D, lower). Scale bar on coronal sections = 2mm, inset = 500um. All sections were obtained 2 weeks after lesioning.

Interestingly, we noted that the reliability of targeting could be functionally evaluated at the time of lesion induction. Under deep anesthesia, sham lesions, as expected, did not produce any noticeable movements (Figure 2B) and Nissl stain at 2 weeks post-operative failed to reveal any changes to the brain parenchyma at the target site (Figure 2D upper panel). However, when the electrolytic charge was administered at the target site, we noticed that stereotyped movements were elicited despite persistent deep anesthesia (no response to deep toe pressure; Figure 2B). With electrolytic charge delivery, we noticed two classes of responses: first were animals that showed mild tremor and/or contraction of the fore limb as well as tremor of the nuchal muscles at the nape (Figure 2B). Upon examination of the tissue in this class of animals we noted that the lesions showed hypercellularity at the lesion site but lacked cystic changes at the core of the lesion (Figure 2D middle panel). The second class of lesions resulted in large amplitude bicycling of the hind limbs, tremor and extension/contraction of the forelimbs, nuchal tremor and contraction along the spine, and enuresis (Figure 2B). Upon examination of the tissue from these animals, cystic degeneration was noted at the core of the lesion site with more extensive inflammatory glial infiltrate around the lesion site (see also Figure 1D-F). Animals with the larger acquired lesions of the second class, as defined by the movements elicited during lesion induction, were used in the subsequently reported studies. This decision was based on the previously observed correlation of percent damage to the bilateral SCPs with penetrance of severity of cerebellar cognitive affective syndrome in patient populations [3], from which it was extrapolated that larger lesions would reflect more robust cerebellar pathology.

### Lesioning of the bilateral cerebellar outflow tracts produces acute, severe dystonia

As described above, the correct stereotactic targeting of the bilateral cerebellar outflow tracts was possible through observation of movements elicited during the delivery of the electrolytic charge. We then observed that lesions of the bilateral cerebellar outflow tracts produced an acute, severe dystonic phenotype. To characterize the dystonia, the mice were observed at 30 mins, 1 hr, 2 hrs, and 24 hrs after surgery. At 30 mins, the lesioned mice were observed to have a fixed, immobile posture, which included a combination of stiff extended limbs, splayed digits, kinked tail, head version, and twisting or extension of the back (Figure 3A, [Video 1](https://1drv.ms/v/s!Auxc2eHbbIGZgcpE2KPDOwpDfVUARA?e=Ih0sRd)). At 1 hr, there was increased movement, with more frequent movements of the limbs and trunk (Figure 3A, [Video 1](https://1drv.ms/v/s!Auxc2eHbbIGZgcpE2KPDOwpDfVUARA?e=Ih0sRd)), though on occasion movement elicited barrel rolling behavior. At 2 hrs, the mice were sometimes able to form a base to stand and take several steps at a time. However, dystonic features including trunk flexion or extension, splayed digits, tail kink, and intermittent flexion and or extension of the limbs continued to be evident (Figure 3A, [Video 1](https://1drv.ms/v/s!Auxc2eHbbIGZgcpE2KPDOwpDfVUARA?e=Ih0sRd)). At 24 hrs, more ambulation was observed, though it was still limited by the dystonic posturing (Fig 3A, [Video 1](https://1drv.ms/v/s!Auxc2eHbbIGZgcpE2KPDOwpDfVUARA?e=Ih0sRd)). To quantify the behavioral dystonia phenotype, we used the previously published dystonia rating scale, which has been used to measure dystonia in mice (Figure 3B) [63]. We found severe dystonia that persisted for up to 24 hrs in the lesion mice, which was significantly worse than the motor phenotype observed in the sham lesion animals (sham n= 5, lesion n=8; two-way mixed ANOVA p<0.0001; Figure 3C). It should be noted that the mice with sham lesions did not display dystonic features at any time after 30 mins; the score of 1 on the dystonia rating scale reflects slowed motor behavior, most likely associated with recovery from surgery (Figure 3C) and the presence of a more subtle lesion that is unavoidable with electrode placement.

**Figure 3.**
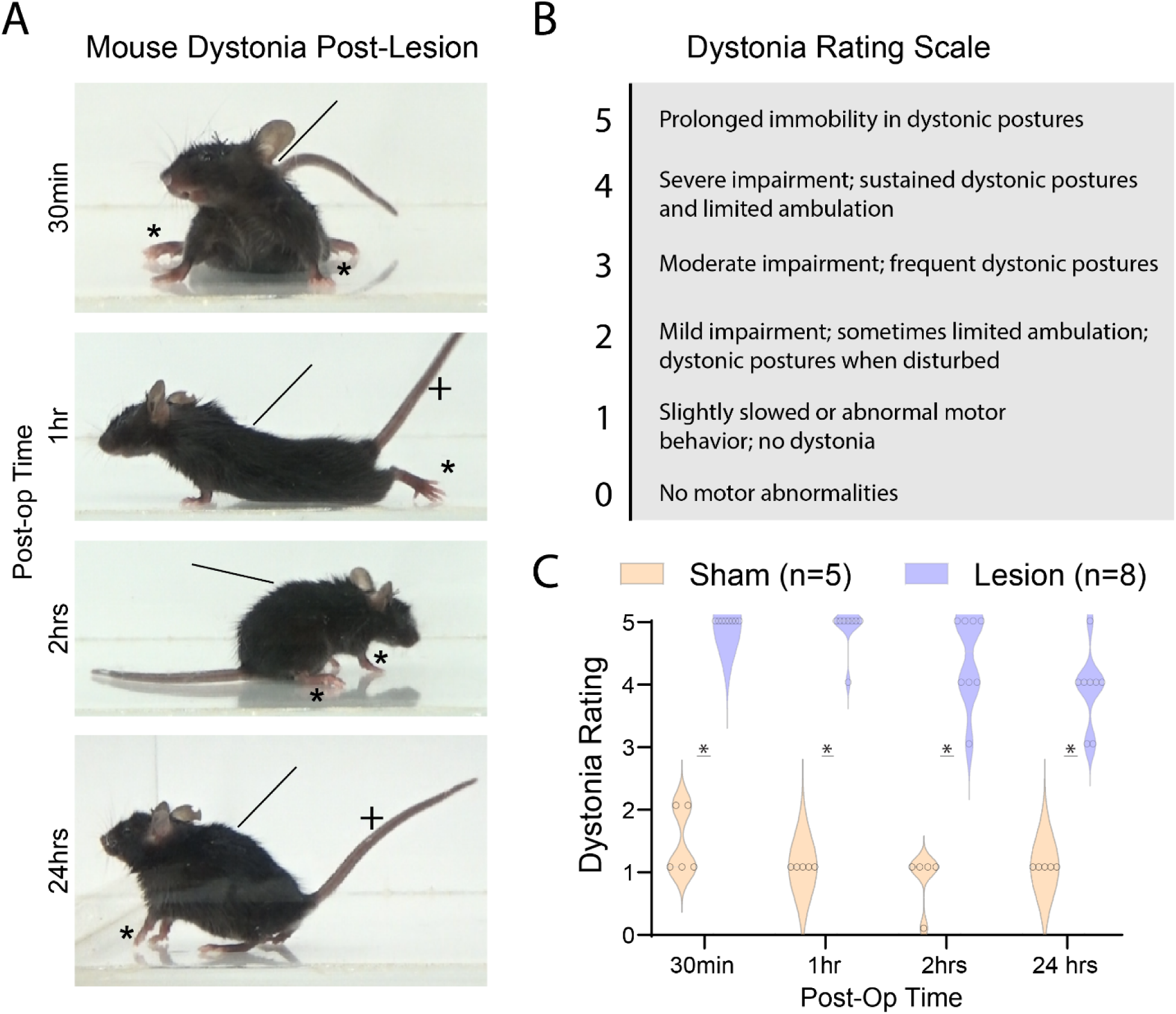
Cerebellar outflow tract lesions lead to acute, severe dystonia. Lesioning of the cerebellar outflow tracts induces an acute severe dystonia apparent as the animal recovers from anesthesia. At 30 mins, the animals display a fixed posture with version of the neck, stiff limbs, and splayed digits (A, 30 min); at 1 hour, there is sometimes improvement, and some steps taken (A, 1 hr); at 2 hours the mice are more often able to perform limited ambulation (A, 2 hrs); by 24 hrs, there is still frequent dystonic posturing but limited ambulation is apparent (A, 24hrs). Asterisks indicate limb/digit dystonia, lines indicate cervical/trunkal dystonia, pluses indicate tail dystonia (A). The dystonia rating scale is used to numerically define these phenotypic observations (B) [63]. Compared to sham lesions (n=5), significantly higher scores on the scale were noted in lesion mice (n=8; two-way mixed method ANOVA p<0.0001) (C).

### Bilateral lesions of the cerebellar outflow tracts do not disrupt Purkinje cell function

Previous mouse models of dystonia mediated by or including cerebellar dysfunction have involved disruption of Purkinje cell function[9, 14, 25, 26, 31, 47, 49, 63, 83, 87]. While several of these studies have also implicated cerebellar nuclear, and thus cerebellar “outflow”, dysfunction in dystonia, they have not delineated whether mechanical disruption of the cerebellar outflow tracts directly leads to dystonic phenotypes and whether disrupted cerebellar nuclear neurotransmission leads to recurrent dysfunction in Purkinje cell firing properties. To understand how cerebellar outflow lesions affect Purkinje cell firing, we performed lesion or sham surgeries on mice and implanted a cannula posterior-lateral to the craniotomy site overlaying the right cerebellar hemisphere (see methods) to accommodate post-operative recording of Purkinje cells. Due to potential persistent effects from anesthesia, we performed awake recordings at 48 hrs, using a protocol that was previously developed in our lab[10, 81].

At 48 hrs post-operative, the lesion mice had improved, but persistent dystonia compared to the sham lesion mice remained (sham n=5, lesion n=8 unpaired t-test p<0.0001; Figure 4A). To characterize the firing properties of Purkinje cells (Figure 4B), we examined several well-established parameters of Purkinje cell function, including firing rate, the coefficient of variance (CV) of the interspike interval, and the spike-to-spike variability (CV2) of both Purkinje cell simple spikes and Purkinje cell complex spikes. Interestingly, we did not note any differences in the simple spike firing properties between lesioned and sham lesioned mice (Sham: 11 cells from 3 mice, Lesion: 12 cells from 3 mice. No significant difference via unpaired t-test; Figure 4C upper). Similarly, the complex spike firing properties were also unchanged between the two cohorts of mice (Sham: 11 cells from 3 mice, Lesion: 12 cells from 3 mice. No significant difference via unpaired t-test; Figure 4C lower). These data indicate that the SCP lesions have little to no lasting functional impact on Purkinje cell activity, which could in theory occur indirectly through feedback loops or directly through retrograde effects from the site of the lesion. Therefore, the dystonia we report here is not mediated by Purkinje cell dysfunction.

**Figure 4.**
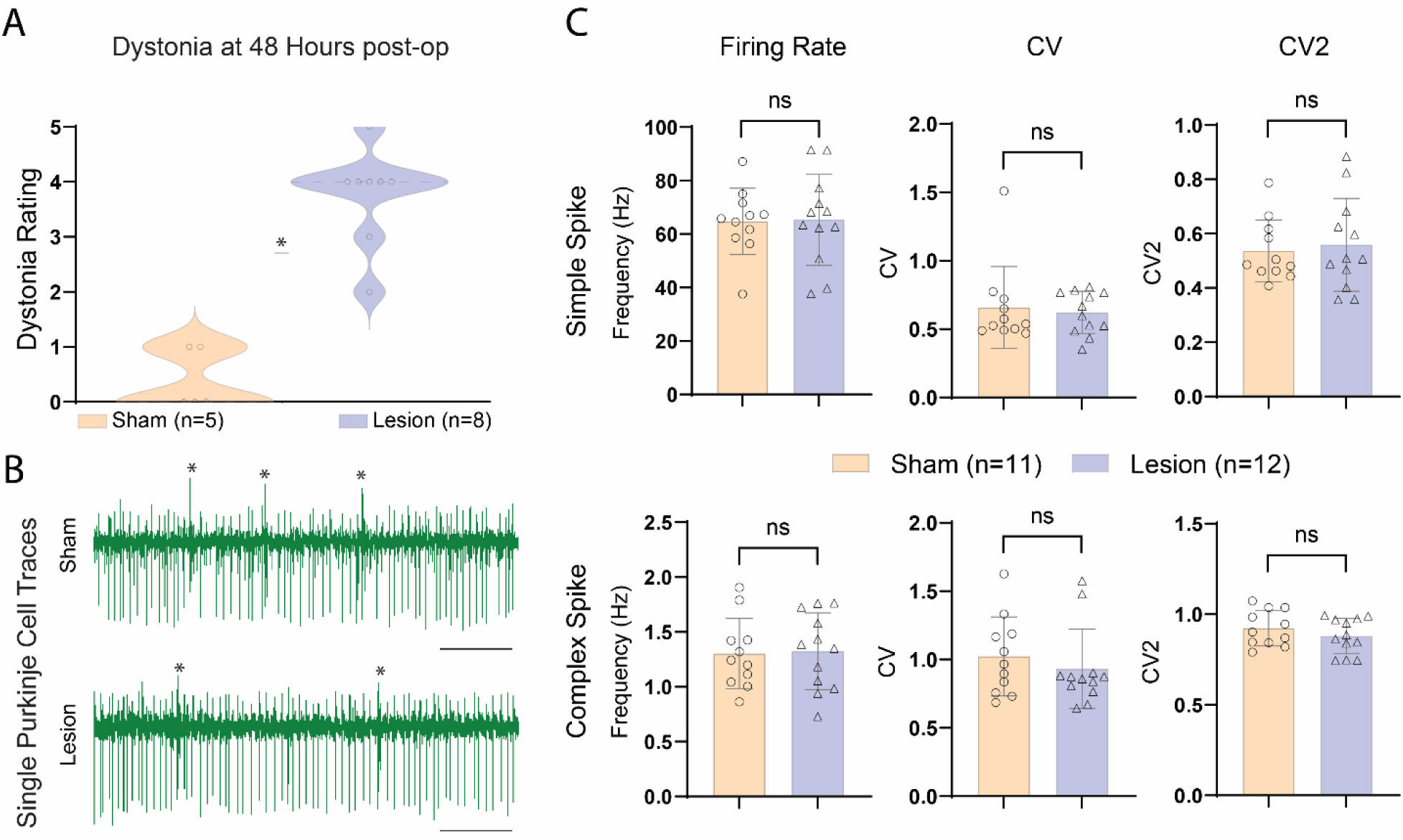
Lesioning of the cerebellar outflow tract does not alter Purkinje cell firing activity. To diminsh the potential effect of anethesia and mechanical damage from craniotomy, Purkinje cell (PC) recordings were performed at 48 hrs, at which time lesion mice continued to have significantly higher dystonia rating scale scores compared to sham lesion animals (unpaired t-test p<0.0001; A). Example traces from single cell recordings of Purkinje cells in a sham lesion mouse (B, upper) and lesion animal (B, lower). Asterisks indicate complex spikes, which were used to confirm that the recording was from a Purkinje cell, scale = 200 ms (B). Purkinje cell simple spike firing properties from sham (N=2, n=5) and lesion (N=3, n=6) revealed no differences in firing rate, CV, or CV2 (C, upper). Purkinje cell complex spike firing properties from sham (N=3, n=11) and lesion (N=3, n=12) mice also revealed no differences in firing rate, CV, or CV2 (C, lower).

### DBS of the centrolateral thalamic nucleus, but not the dorsal striatum, alleviates dystonia induced by cerebellar outflow lesions

Investigation of the heterogenous etiologic and phenomenological presentation of dystonia has lead the field to propose the existence of a “dystonia network” encompassing, as primary nodes, the cerebellum, thalamus, somato-motor cortex, and basal ganglia (Figure 5E) [58, 65, 72]. The characterization of the dystonia network has evolved not in small part due to the success of DBS in the treatment of dystonia. In the case of genetic dystonia, the basal ganglia – in particular the globus pallidus interna (GPi) and subthalamic nucleus (STN) – has emerged as a useful therapeutic target in the treatment of dystonia[57]. GPi DBS, which is the most common target, has shown similar efficacy with both high (130Hz) and low (<60Hz) frequency DBS[4, 21]. However basal ganglia targeting consistently shows decreased efficacy for acquired and heredodegenerative etiologies of the condition[29, 52], increasing the urgency to develop new, effective targets.

**Figure 5.**
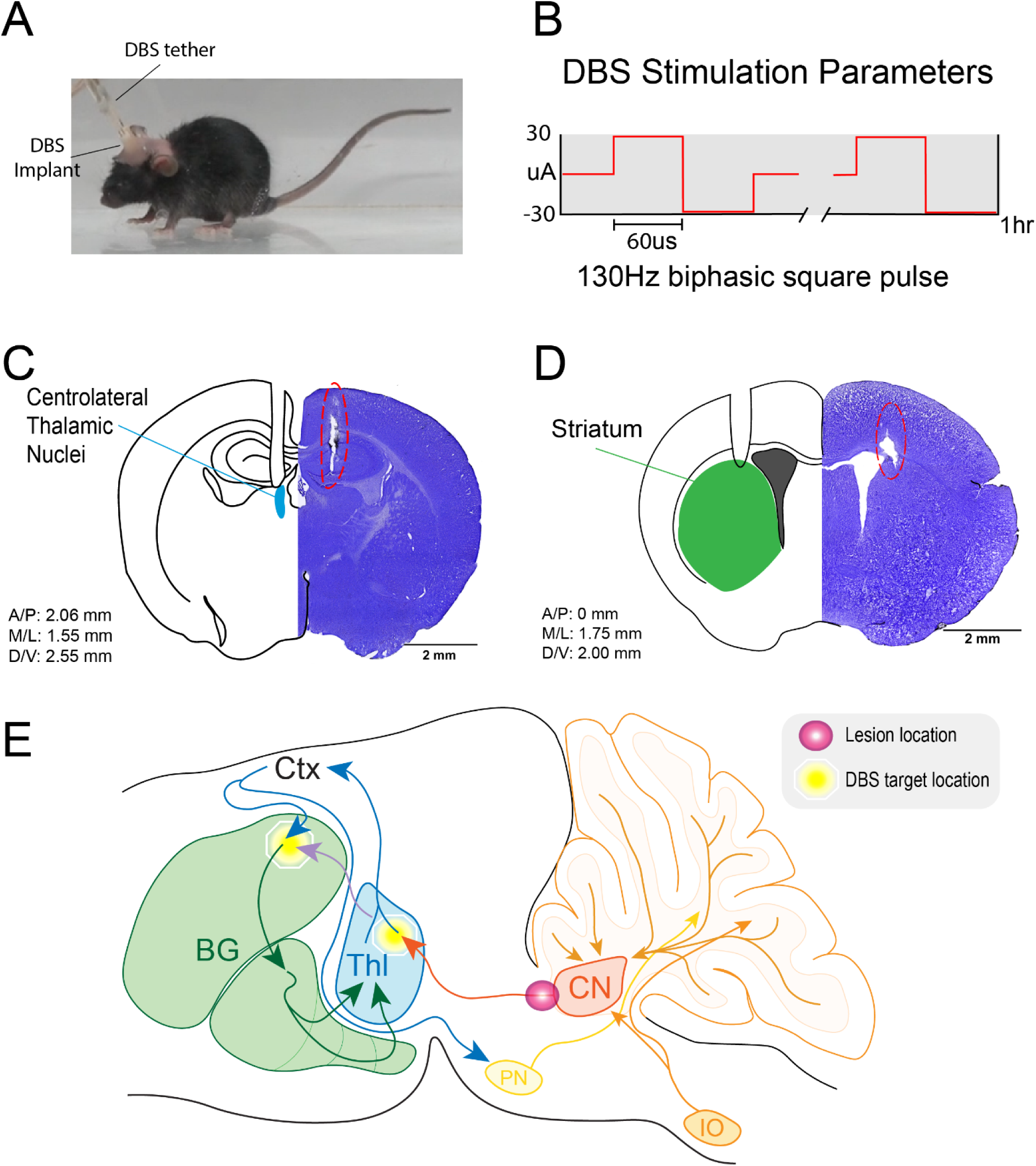
Targeting nodes in the dystonia network with Deep Brain Stimulation (DBS). The DBS implant consists of 2 twisted bipolar tungsten electrodes bonded together at the appropriate distance to target the desired brain region bilaterally (A). DBS stimulation parameters included biphasic square pulses of 30 mA at 130 Hz (B). After DBS stimulation, mice were sacrificed at 2 weeks post-lesion and Nissl stain was used to verify appropriate targeting. Coordinates were obtained from Paxinos and Franklin[62]. Centrolateral thalamic targeting was 2.06 mm posterior from bregma, +/- 1.55 mm along the medio-lateral plane, and 2.55 mm ventral from the brain surface (C). Dorsal striatum was targeted at bregma, +/- 1.75 mm along the mediolateral plane, and 2.00 mm ventral from the brain surface (D). A simplified scheme of the “dystonia network” with lesion location (pink circle) and DBS targets (yellow octagon) indicated. The cerebellar nodes (cerebellar nuclei (CN), pontine nuclei (PN), and inferior olive (IO) are indicated in orange hues, the thalamo-cortical loops (thalamus (Thl), cortex (Ctx)) are indicated in blue, and the basal ganglia (BG) nodes are indicated in green (E). The basal ganglia are divided into the striatum (upper left loop) and deep gray nuclei (lower right loop, consisting of globus pallidus interna, globus pallidus externa, subthalamic nucleus). The short latency disynaptic projection linking the CN to BG is indicated in mauve.

In mice, DBS targeting of the Gpi has remained technologically challenging due to the anatomical and technical constraints of the model system[67]. Furthermore, mouse studies of dystonia mediated by basal ganglia dysfunction have revealed that the dorsal striatum plays an important role in mediating dystonic movements in mice[49, 61] while ablation of the centrolateral thalamic nuclei, which mediates cerebello-striatal signaling, has been shown to alleviate cerebellar etiologies of dystonia[14], implicating this structure in dystonia etiology. Based on these previous observations, we sought to test whether DBS of the dorsal striatum (Figure 5C) or the centrolateral thalamic nucleus (Figure 5D) would be sufficient to alleviate the acute motor features of dystonia that are induced by cerebellar outflow lesions.

Our group has previously successfully used DBS of the cerebellar nuclei to alleviate a variety of movement disorders mediated by cerebellar cortical dysfunction[10, 55, 83]. Several studies have further implicated the centrolateral thalamus, which mediates cerebello-striatal connectivity, in rapid, short latency signaling in dystonia[14, 17]. Furthermore, dystonia resulting from aberrant afferent olivocerebellar signaling can be alleviated by DBS of the centrolateral thalamic nuclei. To test how acute cerebellar outflow-mediated dystonia responds to neuromodulation of these key subcortical nodes of the dystonia network, we fabricated DBS implants consisting of two bipolar tungsten electrodes that could be connected to a pre-programmed pulse generator to deliver 130 Hz biphasic square pulse stimulation of +/- 30 uA to the target site (Figure 5A, B). Stimulation parameters were based on their efficacy in alleviating olivocerebellar-induced dystonia[83]. Target location in either the dorsal striatum or centrolateral thalamic nuclei was confirmed with Nissl stain for all mice used in the experiments (Figure 5C, D).

To test how neuromodulation of nodes in the dystonia network interact with dystonia resulting from lesioning of the cerebellar outflow tracts, we used 130 Hz biphasic DBS (Figure 5D) of the centrothalamic nuclei in mice that had undergone lesioning the day prior. Prior to stimulation, all lesioned mice presented with moderate to severe dystonia including difficulty ambulating, splayed digits, and contraction or twisting of the trunk (Figure 6A left; [Video 2](https://1drv.ms/v/s!Auxc2eHbbIGZgcpE2KPDOwpDfVUARA?e=Ih0sRd)). This was reflected in an average dystonia rating scale score of 3.8 (n=7; Figure 6B, middle). After 1 hr of centrolateral thalamic DBS, all lesioned mice showed significant improvement in dystonic posturing and gait (Figure 6A right, [Video 2](https://1drv.ms/v/s!Auxc2eHbbIGZgcpE2KPDOwpDfVUARA?e=Ih0sRd)), which was reflected in a significant improvement in dystonia rating scale score (post-DBS DRS mean 2.8; paired t-test p<0.0001; Figure 6B middle). This improvement persisted through the immediate post stimulation period before the effect waned in the 60-90 minutes post-stimulation. We next tested whether DBS of dorsal striatum, which is part of a disynaptic circuit with the cerebellar outflow tract and is implicated in dystonia pathogenesis (Figure 5E mauve) [17, 49], would produce similar improvements. After 1 hr of DBS of the dorsal striatum, no phenotypic change was noted in the mice, which was reflected by the absence of change on the dystonia rating scale (Figure 6B, right). We also tested DBS on sham lesioned mice to assay whether there was 1) an independent effect of centrolateral thalamic DBS implantation on dystonia presentation and 2) whether stimulation of the centrolateral thalamic nuclei itself resulted in abnormal motor behavior. We did not note any change in behavior in the sham lesioned animals after 1 hr of centrolateral thalamic DBS, which was reflected in absence of change in the dystonia rating scale (Figure 6B left). As noted previously, sham mice scored a 1 on the scale due to expected post-operative slowness of movement, although no dystonic features were noted. Thus, 1 hr of centrolateral thalamic nuclear stimulation improved dystonic phenotypes in mice, whereas dorsal striatal stimulation did not demonstrate any improvement. All mice in the described experiments underwent a single 1 hr session of DBS.

**Figure 6.**
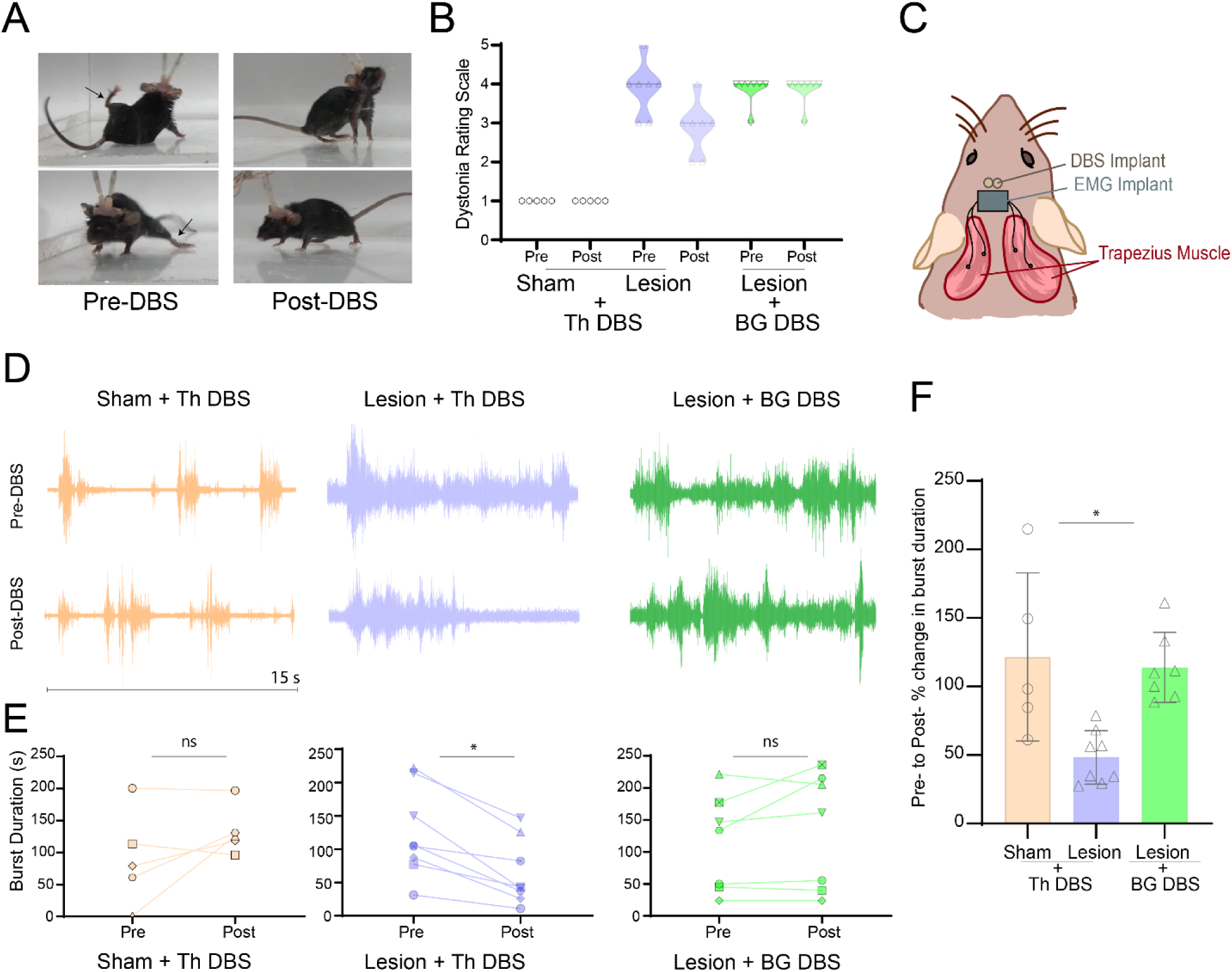
Outcomes of Deep Brain Stimualtion (DBS). Prior to centrothalamic DBS, at 24 hrs post-lesion, mice have ongoing severe dystonia with stiff, splayed limbs (arrows) and trunk contraction (A, left). After 24 hrs of centrothalamic DBS, there is improvement in gait and improvement of tone in the trunk and extremities (A, right). Using the dystonia rating scale (DRS), we note no change in movements pre to post DBS with stimulation of centrothalamic nuclei in sham animals (B; left) or in lesion animals with dorsal striatal DBS (B, right). However, all lesion animals with centrothalamic DBS demonstrated improvement on the DRS, leading to significant group effect (ratio paired t-test p<0.0001). To determine whether muscle contraction was affected by DBS in the three conditions, we used nuchal EMG, with 2 silver wire electrodes inserted into each the left and right trapezius (C). EMG yielded raw burst traces as indicated in (D). Example 15 second traces from an individual sham lesion mouse with centrolateral DBS pre and post stimulation (beige), an individual lesion mouse with centrolateral thalamic DBS pre and post stimulation (blue), and an individual lesion mouse with dorsal striatal DBS pre and post stimulation (green) are shown (D). To evaluate how muscle contraction was affected in individual animals, we calculated total burst duration (a proxy for muscle contraction) over the 5 mintues prior to stimulation and the five minutes just after DBS was switched off in individual animals. We did not find a significant difference using a paired t-test in sham lesion with centrolateral thalamic DBS (n=5, paired t-test p=0.1735; E, left) or lesion animals with dorsal striatal DBS (n=7, paired t-test p=0.2842; E, right). With centrolateral thalamic stimulation in lesion animals, we noted a significant decrease in burst duration from the 5 mins pre DBS to post DBS (n=8, paired t-test p=0.0013; E, center). To compare between conditions, percent change in burst duration after 1 hr of stimulation was examined, revealing a significant percent decrease in the lesion mice with centrolateral thalamic DBS compared to the other two conditions (Brown-Forsythe p=0.0309 and Welch’s p=0.0016 ANOVA tests; F).

### Behavioral improvements after centrolateral nucleus stimulation correspond to decreased abnormal muscle contractions in dystonic lesioned mice

To functionally confirm the alleviation of dystonia with DBS stimulation, we used nuchal electromyography (EMG) to evaluate how muscle burst duration responded to DBS. We chose nuchal EMG as 1) it has previously been shown to reflect the dystonia evident in cerebellar etiologies of dystonia in mice[48, 68]; 2) cervical dystonia is an important dystonic presentation in patients with cerebellar lesions[20]; and 3) it is a technically less challenging surgical intervention for the mice than limb EMG. Nuchal EMG involves the implantation of two electrodes directly into each of the left and right trapezius in the mouse (4 electrodes implanted into the muscle in total) and a ground wire inserted into the subcutaneous tissue overlaying the muscle (Figure 6C). Examples of raw traces from implants are provided in Figure 6D, showing examples of burst activity that reflects muscle contractions. Given the variability in the insertion location, possible mild muscle damage during surgical exposure, and interindividual variability, it was difficult to compare contractions between individuals. We therefore compared “before” and “after” DBS in each mouse. With this measure, similar to what we observed with the phenotypic measures, we found that there was a significant decrease in burst duration after 1 hr of centrolateral thalamic DBS (paired t-test, df=7 pairs, p=0.0013), but no change seen in burst duration after 1 hr of centrolateral thalamic DBS in animals with sham lesions (paired t-test, df=4, p=0.1735) nor in lesioned, dystonic animals after 1 hr of dorsal striatal DBS (paired t-test, df=7, p=0.2842; Figure 6E). To compare the response to DBS between the three groups, we examined the percent change in trapezius muscle burst duration after one hour of DBS in each condition. We found that there was a significant difference in percent change in the lesioned animals with centrolateral thalamic DBS compared to the other groups (Brown-Forsythe p=0.0309 and Welch’s p=0.0016 ANOVA tests; Figure 6F).

## Discussion

We developed an electrolytic lesion paradigm to investigate the effect of acquired injury of the cerebellar outflow tract on dystonia pathogenesis and the interaction of this injury with nodes in the dystonia network during therapy. We found that focal lesions of the cerebellar outflow tract, which interrupt efferent signaling from the cerebellar cortex to the thalamus, cerebral cortex, and basal ganglia, result in an acute severe phenotype that is consistent with the key features of dystonia in mice. Investigation of Purkinje cell function after lesion induction revealed an absence of recurrent dysfunction in Purkinje cells, indicating that disruption of efferent neurotransmission at the level of the cerebellar nuclei was sufficient to induce dystonia without altering the Purkinje cells. Furthermore, we found that DBS of a downstream first order node of the network, the centrolateral thalamic nucleus, alleviated the dystonia but stimulation of a short latency second order node, the dorsal striatum, did not achieve such an effect. Together, we found that disruption of cerebellar outflow is sufficient to induce an acute, severe dystonia that shows rapid response to centrolateral thalamic, but not dorsal striatal, DBS.

### Induced damage of the cerebellum leads to dystonic motor behaviors in mice

Acquired injury to various components of a broad “dystonia network” have been reported to cause dystonia, with the cerebellum and basal ganglia emerging as critical nodes in this network[18, 59]. Previously, excitotoxic lesions of the cerebellar hemispheres or vermis, using kainic acid, have shown that injury to cerebellar components of the dystonia network can lead to dystonia[27, 59, 63, 79], while pharmacologic manipulations of the cerebellum in mouse models predisposed to dystonia also elicit dystonia[14]. Non-pharmacological lesioning of the cerebellar outflow tracts has not, to our knowledge, been reported in mice in the context of dystonia, although models of injury to the medial cerebellar nuclei and vermis have been reported in rats, using electrolytic charge delivery and mechanical suction, respectively[1, 8, 33, 34]. These studies do not report on dystonia, although the study on medial cerebellar nuclei lesions reported difficulty with ambulation in the early post-operative days[1].

Here, we induced acute onset severe dystonia after bilateral lesioning of the SCPs. The observation of stiffened limbs, contraction of the cervical and paraspinal muscles, and splayed digits phenocopy the movement disorder as seen in the clinical diagnosis of dystonia. Furthermore, based on the dystonia rating scale[9, 14, 63, 83], the overall phenotype is consistent with a severe dystonia. Importantly, in patients, qualitative evaluation of phenotype-based clinical characterization of movements and postures is a critical determinant of disease presentation and treatment response[12]. Finally, it is important to note that in etiologically complex movement disorders arising from acquired injury, such as cerebral palsy, it is often clinically difficult to distinguish between “competing” diagnoses such as spasticity and dystonia[22, 53]; paramount is arriving at a treatment that improves quality of life.

In human patients with lesion burden similar to what we have induced in this study, dystonia is not necessarily the most prominent feature of the clinical presentation[3]. Rather, these patients present with ataxia as the predominant movement disorder and with mutism as the most important “non-motor” symptom. However, the presentation can include dystonia and the characterization of the mutism is still an area of active consideration[41, 84]. Furthermore, the trajectory of the resulting constellation of neurologic deficits, coined “cerebellar mutism syndrome”, starts with a severe early presentation that wanes over an intermediate time frame (3-6 months in humans), similar to what is seen in our study. The similar trajectory (acute onset with subsequent waning) and severity (severe with gradual reduction in scope of symptoms) highlight the importance of understanding how the network downstream of the cerebellar outflow tract responds and adapts to the insult. As a result, ongoing characterization of the symptoms of the cerebellar outflow lesion model and network response is of great importance. Whether small lesions, lesions in different locations (more proximal/distal to the nuclei; closer to the lateral nuclei, etc.) or lesions at different ages would induce a different constellation of symptoms will also be important to test. Our observations highlight the importance of understanding and implementing principles derived from network studies to guide node specific DBS in the treatment of disease.

### Therapeutic modulation of acute cerebellar outflow in dystonia

DBS of the globus pallidus interna is a key therapeutic intervention in primary, usually genetic, dystonia[36, 39]. Thalamic DBS has been used with efficacy in dystonia that does not respond to GPi, generally in the case of acquired dystonia[51, 86]. Here, we report that the acute severe dystonia resulting from cerebellar outflow lesions shows alleviation in less than 1 hr with DBS of the centrolateral thalamic nuclei, but no improvement after 1 hr of dorsal striatum stimulation. This reduction in dystonia is qualitatively apparent on visual examination of the mice, quantitatively measured based on the dystonia rating scale and correlates with a significant decrease in contraction duration of the trapezius muscle. The absence of efficacy of dorsal striatal DBS in the alleviation of dystonia comes as a surprise given the efficacy of basal ganglia DBS in primary dystonias. Reasons for our observed lack of efficacy include the relatively short duration of stimulation in our paradigm, as well as the targeting of the striatum rather than the GPi, which, as mentioned, is not feasible in rodents as this structure is not fully conserved.

An intriguing aspect of our observations is the short latency to response of the dystonia with DBS. As opposed to DBS of the STN in Parkinson’s or the ventral striatum in OCD, in which symptoms can show rapid, nearly instantaneous response, DBS of the GPi leads to alleviation of dystonia on the far more protracted scale of days to weeks, or in some cases months[69]. Previously, cerebello-cortical and basal ganglia cortical loops were thought to interact through slow top-down cortical control[54]. More recently, fast disynaptic connectivity between the cerebellum and basal ganglia, in particular via a single node connecting the cerebellar nuclei and the striatum, was shown in mice[17]. Our results build upon this observation and indicate that perhaps this short latency circuit may be responsible for both 1) the rapid onset of symptom presentation in acquired cerebellar dystonia and 2) the relatively rapid DBS response noted in our study. An interesting paradox here is that DBS of the centrolateral thalamus, a first order node from the SCP, and not the striatum, a second order node in the same pathway, is better at reducing dystonic symptoms. This strongly suggests that even within a single pathway, the exact node chosen for DBS targeting can have a dramatic impact on the therapeutic outcomes. Increasingly, DBS exploiting the key position of the cerebello-thalamic link has been undertaken in clinical studies: in dystonia, SCP and cerebellar outflow DBS have shown promise[13, 38]; and early results indicate that centrolateral thalamic DBS has efficacy in recovery from moderate to severe traumatic brain injury[71]. Intriguingly, recent work has demonstrated that the effect of cerebellar stimulation on stroke recovery in mice depends on the integrity of the striatum[16], directly linking the dorsolateral striatum, cortex, cerebellar projections, and their unifying node, the thalamus, which is the target of modulation in the present study. Given the key position of the cerebellothalamic projection in the sensorimotor machinery and the increasingly evident impact that neuromodulation of this link has on disorders of sensorimotor integration, using the present model to understand how the cerebellum modulates neuroplasticity presents an exciting avenue.

In particular, expansion of interrogation of the dystonia network beyond the short latency cerebello-thalamo-striatal circuit undertaken in the present study should focus on plasticity of both extracerebellar (GPi) nodes and the preserved cerebellar nuclei; DBS of these two nodes was precluded in the current study due to 1) the inaccessible anatomic location of the GPi in mice and 2) the proximity of the lesions to the cerebellar nuclei, which caused our trial experiments for cerebellar nuclear targeting to fail due to subject morbidity and mortality. Whether modulation of the GPi and cerebellar nuclei operate with short- or long-term latency to improve dystonia could reveal how components of the dystonia network interact in disease and compensation after injury.

## Conclusion

This study highlights the use of brain network structure and function in understanding the genesis of dystonia and uses interregional connectivity to inspire a targeted neuromodulation approach for symptom management. The conserved intrinsic and first order cerebellar circuitry allow for this lesion-based model of dystonia to uncover insights on how the cerebellum and its associated networks respond to neuromodulation and recover from acquired injury.

## Acknowledgements

This work was supported by Baylor College of Medicine, Texas Children’s Hospital, the National Institute of Neurological Disorders and Stroke (RVS: R01NS100874, R01NS119301, and R01NS127435; JSG: K08NS121600), Eunice Kennedy Shriver National Institute of Child Health and Human Development of the National Institutes of Health under Award Number P50HD103555 for use of the Cell and Tissue Pathogenesis Core and *In Situ* Hybridization Core (the BCM IDDRC). The content is solely the responsibility of the authors and does not necessarily represent the official views of the National Center for Research Resources or the National Institutes of Health. The authors have no relevant financial or other conflicts of interest to declare.

## Contributions

Megan X. Nguyen, Amanda M Brown, Roy V Sillitoe, and Jason S Gill authors contributed to the study conception and design. Material preparation, data collection and analysis were performed by MXN, AMB, Tao Lin, and JSG. The first draft of the manuscript was written by JSG and all authors commented on previous versions of the manuscript. All authors read and approved the final manuscript.

## Ethics

Animal experimentation: Mice were housed in an AAALAS-certified animal facility. All procedures to maintain and use these mice were approved by the Institutional Animal Care and Use Committee for Baylor College of Medicine (Animal protocol number AN-5996).

